# The Relationship between Resilient Mangroves and Fish Populations in the Largest Marine Reserve in Belize: A Case for Conservation

**DOI:** 10.1101/719757

**Authors:** Chetwynd Osborne, Leandra Cho-Ricketts, Jané Salazar

**Affiliations:** Environmental Research Institute, University of Belize, Belmopan, Belize

## Abstract

Mangrove forests are one of the most bio-diverse and productive wetland environments on earth. However, these unique tropical forest environments that occupy coastal areas are among the most threatened habitats globally. Approximately one-third of the world’s mangrove forests in tropical and subtropical coastlines have been lost to logging, conversion of land for agriculture and mariculture, and degradation due to general pollution over the past 50 years. The large population of mangroves occupying the Turneffe Atoll area in Belize face growing anthropogenic threats such as permanent clearing of land for housing and infrastructural development as well as pollution and natural factors (climate change). Despite these threats, mangroves have high resilience due to their fast rates of microbial decomposition and nutrient flux, large storage of nutrients below-ground, numerous feedbacks, keystone species redundancy and self-design, and highly efficient biotic controls. Given the few formal studies done to evaluate mangrove resilience at Turneffe Atoll, the purpose of this study was to evaluate mangrove resilience and nursery functions in the Turneffe Atoll Marine Reserve. Mangrove fish abundance, forest structure, and prop root length was assessed by means of visual census and point centred quarter method (PCQM) for 11 sites that span across conservation and general use zones. This study found that resilient mangroves (lower vulnerability ranks, higher standing biomass, and higher fish biomass and abundance) exist in conservation and general use zones and warrant the need for improved mangrove conservation measures. Some conservation measures include: greater reduction of non-climate stressors, establishment of more mangrove conservation areas, and implementation of more outreach and educational programmes.

## Introduction

Mangroves are classified as small evergreen trees that thrive in intertidal zones of estuaries, lagoons, river deltas, and dominate subtropical and tropical coastal systems [1–5]. The productive nature and location of mangroves in nearshore, warm coastal waters, make them increasingly valuable targets for farming, mariculture, and recreation. These activities fundamentally alter the physicochemical nature of the habitat, affecting animals such as shrimps, crabs, and fishes that depend on these ecosystems for food and shelter [6,7]. Habitat destruction of coastal mangrove habitats through sea-level rise by means of climate change and deforestation by means of anthropogenic activities can alter niche dynamics in these communities [8,9]. Mangrove forests are declining globally due in part to coastal development which has resulted in the loss of one-third of mangrove forests worldwide over the past 50 years [1,3]. Mangrove forests are important for the sustenance of fishes and invertebrates; providing coastal populations with protein sources and supporting livelihoods; shoreline protection against floods, tsunamis, and typhoons; purification of water; absorption of pollutants; offsetting greenhouse gas emissions and sequestering carbon; and provision of nursery habitats for fishes [2,5,8,10–13]. Additionally, these vital ecological goods and services have the potential to improve mangrove resilience to climate change, storm, sea level rise, and anthropogenic activities [8,11].

The most common variable used to illustrate mangrove species zonation patterns is tidal inundation frequency [14,15]. Mangrove species zonation in Belize adheres to the pattern of *Rhizophora* typically encountered near the shoreline and inundated areas, while *Avicennia* are frequent in drier areas that are further inland and have higher soil salinities due to evaporation [4,12]. Classifying mangroves into different types provides critical information about the forest structure and facilitates prioritization of conservation efforts. Mangrove seedlings are individual trees < 1.37 m tall and the presence of mangrove seedlings gives an indication of the level of recruitment in a particular area [16,17]. Saplings are trees with a DBH < 2.5 cm while overstory trees are defined by DBH > 5 cm [18–20]. Relatively high growth rates are characteristic features of mangrove saplings and seedlings and these mangrove types are capable of colonizing newly created intertidal substrates and forest gaps [21]. Dwarf mangroves are trees typically < 3 m tall which are constituted by scrub categories and typically occupy areas where the environmental conditions are extreme and the vegetation is restricted to short mangroves [12]. Medium mangroves are trees approximately 3 – 8 m tall which thrive in locations inland that are densely populated by fringing *R. mangle* which grow in exposed sites along the coast and on cays [12]. Tall mangroves are trees approximately > 7 m tall which are restricted to locations inland and parts that are elevated on the largest cays where the required combination of restricted salinity, high levels of sustained nutrients, and stable substrate exist to sustain this stand of large trees [12]. Standing biomass gives an indication of the quantity of standing organic matter per unit area at a particular time [22]. The quantity of standing biomass stored in a forest is a function of ecosystem age, productivity, exportation strategies, and organic matter allocation [22].

Mangrove ecosystems play a vital role in the Belize Barrier Reef Complex owing to the support provided for animals, from fishes to crustaceans, as well as sustaining high levels of primary production centred around production of leaf-litter, which provides a trophic subsidy for adjacent coastal waters, both near and far [23,24]. Mangrove resilience refers to a measure of the persistence of mangrove ecosystems and of their ability to absorb change and disturbance and still maintain the same relationships between populations [9,11,25,26]. Further, mangroves possess considerable resilience to sea-level fluctuations owing to their ability to actively modify their environment by changing processes of surface elevation, and their migration ability to inland zones over consecutive generations [27]. This sort of migration to landward zones allows the mangrove ecosystem to absorb and recognize the effects of the stress and facilitate the maintenance of its processes, structures, and functions [11,28]. Studies conducted in Bermuda reported that mangrove zones migrated inland where preferred frequency of inundation, depth, and period existed, since these mangroves were incapable of keeping pace with relative sea-level rise [29]. This gives an indication of mangroves’ ability to adapt to changes in sea-level and remain resilient.

Mangroves exhibit high resilience subsequent to disturbances due to their pioneering species ability, high productivity, and regeneration capabilities [26,30]. For instance, mangroves quickly re-grow subsequent to hurricanes (a disaster that Belize is prone to) and floods, but changes in temperature, hydrologic fluxes, topography, and sedimentation affect mangrove forests [30]. Studies conducted in Havana, Cuba, that focused on the effects of permanently increased water levels on mangrove species seedlings revealed that *Rhizophora mangle* seedlings were in a better state when compared other mangrove species (such as *Avicennia germinans* and *Laguncularia racemosa*) and pseudo-mangrove species (such as *Conocarpus erectus*). This was due to *R. mangle* seedlings’ ability to form erect saplings when directly anchored on the base of flooded sites, or lying dead stems or on stumps [26,31]. Although all other species’ saplings are inclined to fall when developing at a greater distance away from the seedling stage, *R. mangle* prop roots promote structural stability and facilitate nutrient exploration and water uptake directly in flooded ground [26,31]. Therefore, different species of mangroves are resilient to varying degrees of flooding due to their regeneration ability under dissimilar flooding regimes.

Mangroves’ complex structure form aquatic vegetation that provides feeding grounds and shelter for small predators and prey [32–38]. For instance, in the Caribbean high abundance and diversity of estuarine, coral reef, and juvenile fishes shelter in mangroves which are structure-rich habitats [38–41]. High abundances of juvenile fishes in mangroves are based on these proposed hypotheses: (a) isolated location of these biotypes from off-shore waters or coral reef, thus less encounters with predators [32,34], (b) copious amounts of food are provided by these biotypes for fishes [34,42], (c) predators efficiency to forage are reduced by the relatively turbid water of the estuaries and bays [33,34], and (d) extensive areas are covered by these biotopes and planktonic fish larvae may be intercepted more effectively [32,34]. These hypotheses support the features of resilient mangroves which provide nursery habitats for juvenile fishes that serve as a protein source for the Belizean population; hence, there is need for further protection of these mangroves at Turneffe Atoll to safeguard this valuable ecosystem service.

Belize still retains large areas of mangroves (especially Turneffe Atoll where two thirds of the atoll’s land area is occupied by mangroves) as compared to many neighbouring countries due to the small population which reduces developmental pressures and the concentration of the population in a single centre (*i.e*., Belize City) [12]. This relatively large area of mangroves should not be treated with complacency, instead action plans should be enacted to further protect these mangroves and prolong the provision of vital ecological goods and services. Mangrove ecosystems are known for their resilience [43] and studies have shown that about 96 hectares of cleared mangrove areas in Belize between the period of 1980 and 2010 have regrown [13]. This sort of regrowth renders the need to better understand the resilient characteristics of mangroves and propose conservation measures to protect this ecosystem and prolong the valuable goods and services provided in terms of maintaining fisheries and supporting biodiversity. The large population of resilient mangroves occupying the Turneffe Atoll area in Belize face growing anthropogenic threats such as permanent clearing of land for housing and infrastructural development as well as pollution and natural factors (climate change) [13]. These threats lead to irreversible losses of mangroves and therefore threatens mangrove resiliency [44]. To date, there have been few formal studies (*e.g*., [45]; [4]) done to evaluate mangrove resilient features in Turneffe Atoll. Therefore, the aim of this research was to evaluate mangrove resilience to support biodiversity composition and nursery functions at Turneffe Atoll. The aim of this research was achieved through the following objectives: (a) to evaluate the resilience of mangrove ecosystems within the Turneffe Atoll, (b) to compare mangrove resilience in conservation and general use zones within the Turneffe Atoll, (c) to evaluate the effect of mangrove prop root structure on fish populations, and (d) to propose necessary conservation measures to further protect mangroves in Turneffe Atoll.

## Materials and methods

### Study area

The study was carried out in Turneffe Atoll (Fig 1), located southeast of Ambergris Caye and Caye Caulker, off the coast of Belize in Central America (17.4382°N, 87.8304°W) [46]. The Turneffe Atoll is made up of many Cayes and most are covered with mangrove forest (covering 74.2 km^2^), while the perimeter of Turneffe Atoll consists of a continuous fringing reef [46]. The sampling design tool for ArcGIS 10.4.1 [47] developed by NOAA’s Biogeography Branch was used to select sample sites that are representative of mangrove distribution (strata) at Turneffe Atoll [48]. Stratified random sampling was done across 11 study sites which span across conservation zones (Long Bogue Conservation Zone [V], Caye Bokel Conservation Area [VI], and Preservation Zone [VII]) and General use zone ([VIII]) at Turneffe Atoll [49].

**Fig 1.**
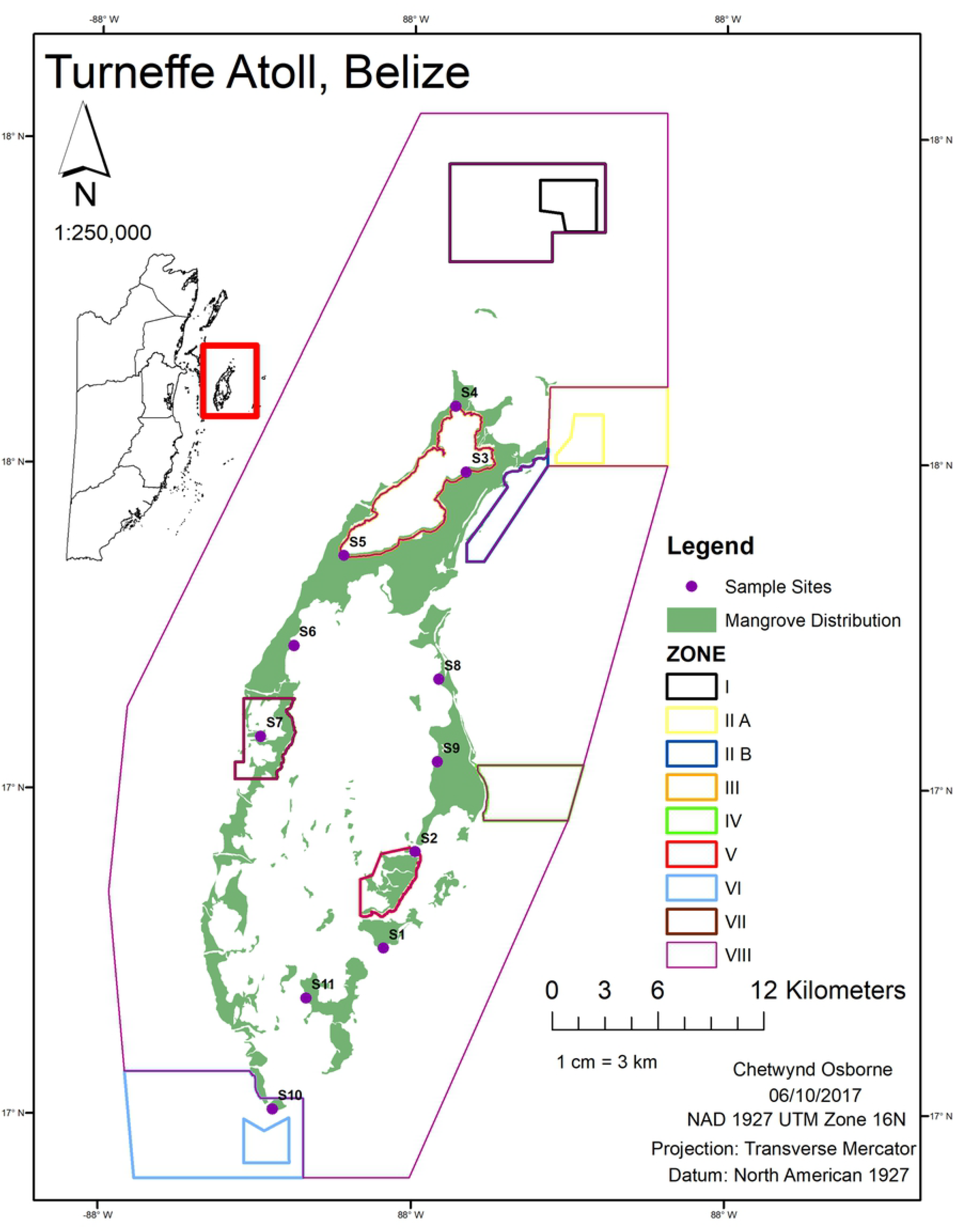
Map of surveyed sites. Zones: I - Maugre Caye Conservation Zone, II A - Dog Flea Conservation Zone, II B - Cockroach – Grassy Caye Special Management Area, III - Vincent’s Lagoon Special Management Area, IV - Blackbird Caye Conservation Zone, V - Long Bogue Conservation Zone, VI - Caye Bokel Conservation Area, VII - Preservation Zone, VIII - General Use Zone.

### Resilience of mangrove ecosystems

The point centred quarter method (PCQM) suggested by [22] and also utilized by [19] was used to assess mangrove forest structure at each mangrove site. PCQM provided some community structure data about mangroves in conservation and general use zones. This sort of measurement provided an indication of mangrove composition to aid with the evaluation of how mangroves were performing within conservation zones versus the general use zone. Four quarters were defined at each sampling point where the transect line and a perpendicular line cross and measurements such as distance from the sampling point to the midpoint of the nearest tree (d, in metres), species, diameter at breast height (DBH, in centimetres), and height (h, in metres) was taken from the tree closest to the sampling point [19,22].

Level of mangrove recruitment, condition, and basal area (m^2^/ha) at each site were determined by rank criteria presented in Table 1, where 5 was high vulnerability and 1 was low, and results were averaged to give an overall mangrove vulnerability rank: 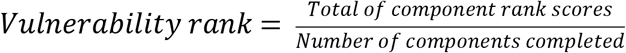 [17,50,51]. Measurements of *R. mangle* prop root in the upper intertidal zone of each belt transect was measured at mangrove prop roots leading edge at a water depth of ca 0.5 m and sometimes measurements were conducted at or near the sediment-water interface close to shore [39].

**Table 1.**
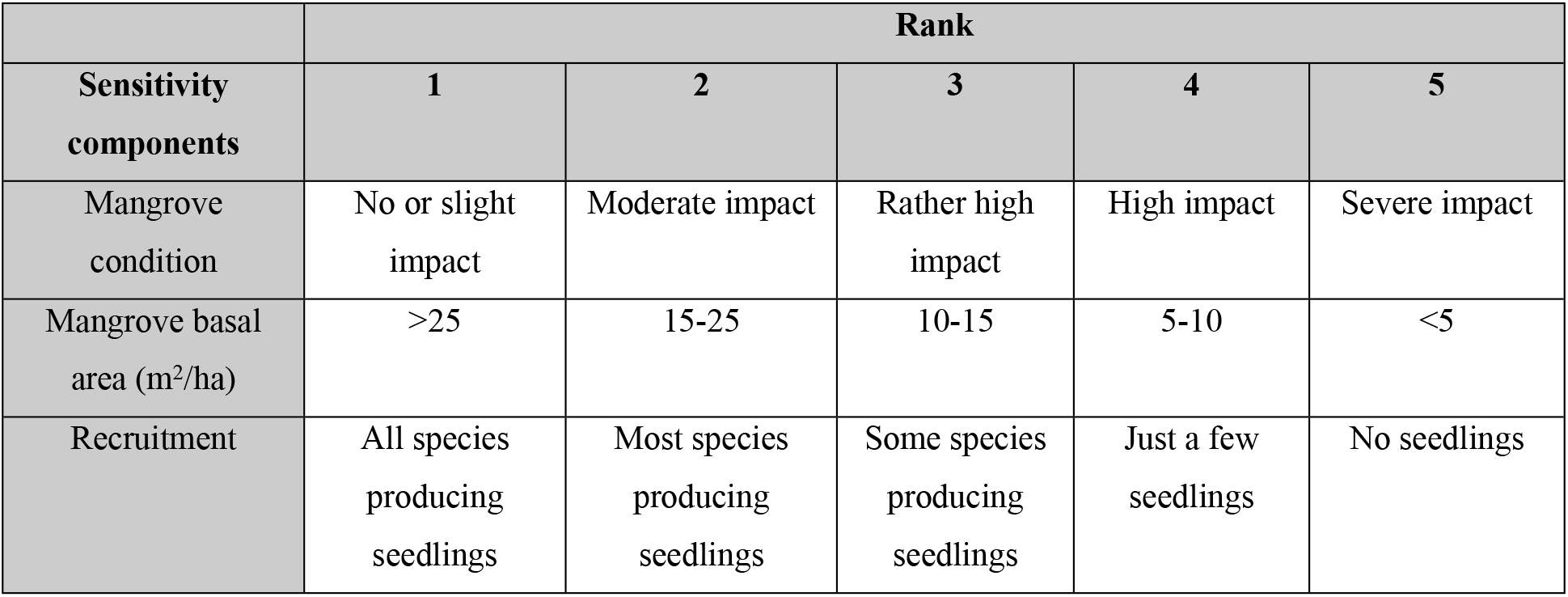
Ranking criteria for mangrove vulnerability assessment results [17].

| Sensitivity components | Rank |  |  |  |  |
| --- | --- | --- | --- | --- | --- |
|  | 1 | 2 | 3 | 4 | 5 |
| Mangrove condition | No or slight impact | Moderate impact | Rather high impact | High impact | Severe impact |
| Mangrove basal area (m<sup>2</sup>/ha) | >25 | 15-25 | 10-15 | 5-10 | <5 |
| Recruitment | All species producing seedlings | Most species producing seedlings | Some species producing seedlings | Just a few seedlings | No seedlings |

### Effect of mangrove prop root structure on fish populations

Visual censuses were used to evaluate fish abundance within conservation zones as compared to general use zone and these censuses were done during the daytime period (08.00–15.00h) to ensure consistency in fish presence and activity [7,34,52–54]. Fish biomass and density recorded across surveyed sites aided in the determination of effects of mangrove prop root structure on fish populations within conservation zones versus the general use zone. The visual census technique [34,55] was used to estimate fish abundance and body length of fish species in mangrove prop-roots at each site. To avoid startling any fish within the two 3 x 30 m belt transects, the observers entered the water at least 20 m from each site [55]. The census was conducted by swimming slowly along the belt transects and the best estimation by eye of abundance and body length of fish species was recorded [34,53]. Size classes of 5 cm were used for total body length estimation and length estimation was guided by graduation marks on the underwater slates that were used for data recording [34]. Water clarity was good for visual censuses across all sites except for site 5 which had poor water clarity. Site five (considered an outlier) was removed for statistical analysis with fish data.

### Statistical analyses

All data were analyzed using the statistical programme R version 3.1.0 [56], at an acceptable α-level of 0.05 (95% confidence level). The PCQM results were used to calculate tree structural variables such as density per centre point 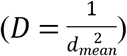, where *D* = stem density in m^−2^ and *d*_*mean*_ = 2 mean distance for all trees on a transect; mean DBH; mean height; basal area per tree 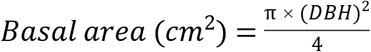, where π = 3.142; and standing biomass per tree (*Biomass (g)* = *b*[(*DBH*)^2^(*height*)]^*m*^), where m and b were constants of 0.8557 and 125.9571, respectively [4,19,22,51]. These densities, means, basal area, and standing biomass provided an indication of mangrove forest structure and the health of mangrove stands. The non-parametric Kruskal-Wallis test (data did not follow normal distribution) was used to test if there were any significant differences in mangrove height and DBH between sites in conservation and general use zones [57,58]. Height and DBH readings for sites in conservation and general use zones were compared using the Mann-Whitney test to evaluate any differences in forest structure between these locations, since they were all a part of Turneffe Atoll [57]. Non-parametric Spearman rank correlation analyses were used to evaluate relationships between *R. mangle* prop root surface area and *R. mangle* standing biomass, between fish abundance and mangrove vulnerability rank, and between fish abundance and mangrove prop root surface area [7].

Fish species were grouped into families, and estimated fish abundance for the four most dominant families among study sites were subjected to *t*-test to determine statistical differences between study sites [34]. The Shannon-Weiner diversity index was used to evaluate fish species diversity between study sites. Size estimates and published length-weight relationships available at FishBase [59] were used to estimate fish biomass [60–64]. Multiple linear regression analysis [65,66] was used to evaluate relationship between *R. mangle* standing biomass and prop root surface area with fish abundance.

## Results

### Mangrove ecosystems resilience within Turneffe Atoll

Based on the 11 sites surveyed across Turneffe Atoll (Fig 1), two species of mangroves (*Rhizophora mangle and Avicennia germinans*) were found across all sites and mangrove ecosystems were dominated by *R. mangle* (Fig 2) overall. However, three sites (S1: general use zone, in central lagoon; S2: conservation zone V and S3: general use zone, close to Vincent’s lagoon special management area) were dominated by *A. germinans* (Fig 2). *Laguncularia racemosa*, one of the least common true mangroves found in Belize [12], was absent from all surveyed sites. Based on the ranking criteria presented in Table 1, all sites had some components of vulnerability (Table 2 and Fig 3). Sites S1, S2, S5, S8, S9, and S10 had ranks of 1 – 2, indicating current resilience for these mangrove areas, while sites S3, S4, S6, S7, and S11 had ranks of 2 – 4, which indicated some moderate vulnerability for these mangrove areas [17]. No statistically significant relationship (*p* = 0.13, Spearman’s correlation) existed between fish abundance and vulnerability rank, indicating that fish abundance was not strongly dependent on the vulnerability state of mangroves. Taking into consideration all sites in conservation and general use zones, higher forest structure measurements in terms of height, DBH, and standing biomass were recorded for general use zone sites (Fig 4), suggesting more healthy and older mangrove stands among these sites. Statistically significant difference (Kruskal-Wallis test, *p* < 0.01) in height and DBH existed among *R. mangle* and *A. germinans*, indicating that mangrove trees of the same species differed in heights and DBH among the sites.

**Table 2.** Vulnerability assessment ranking results for sites in conservation and general use zones. The last row gives averaged rank results of the overall vulnerability rank for each site (shown in bold).

| Sensitivity components | Conservation Zone |  |  | General Use Zone |  |  |  |  |  |  |  |
| --- | --- | --- | --- | --- | --- | --- | --- | --- | --- | --- | --- |
|  | S2 | S7 | S10 | S1 | S3 | S4 | S5 | S6 | S8 | S9 | S11 |
| Mangrove condition* | 1 | 1 | 1 | 1 | 2 | 2 | 1 | 2 | 1 | 2 | 1 |
| Mangrove basal area (m<sup>2</sup>/ha)* | 2 | 5 | 2 | 2 | 3 | 5 | 2 | 4 | 2 | 3 | 5 |
| Recruitment* | 1 | 1 | 1 | 1 | 2 | 1 | 1 | 2 | 1 | 1 | 1 |
| Total | 4 | 7 | 4 | 4 | 7 | 8 | 4 | 8 | 4 | 6 | 7 |
| Number of components | 3 | 3 | 3 | 3 | 3 | 3 | 3 | 3 | 3 | 3 | 3 |
| Vulnerability rank | **1.3** | **2.3** | **1.3** | **1.3** | **2.3** | **2.7** | **1.3** | **2.7** | **1.3** | **2** | **2.3** |

\*Values based on ranking criteria presented in Table 1.

**Fig 2.**
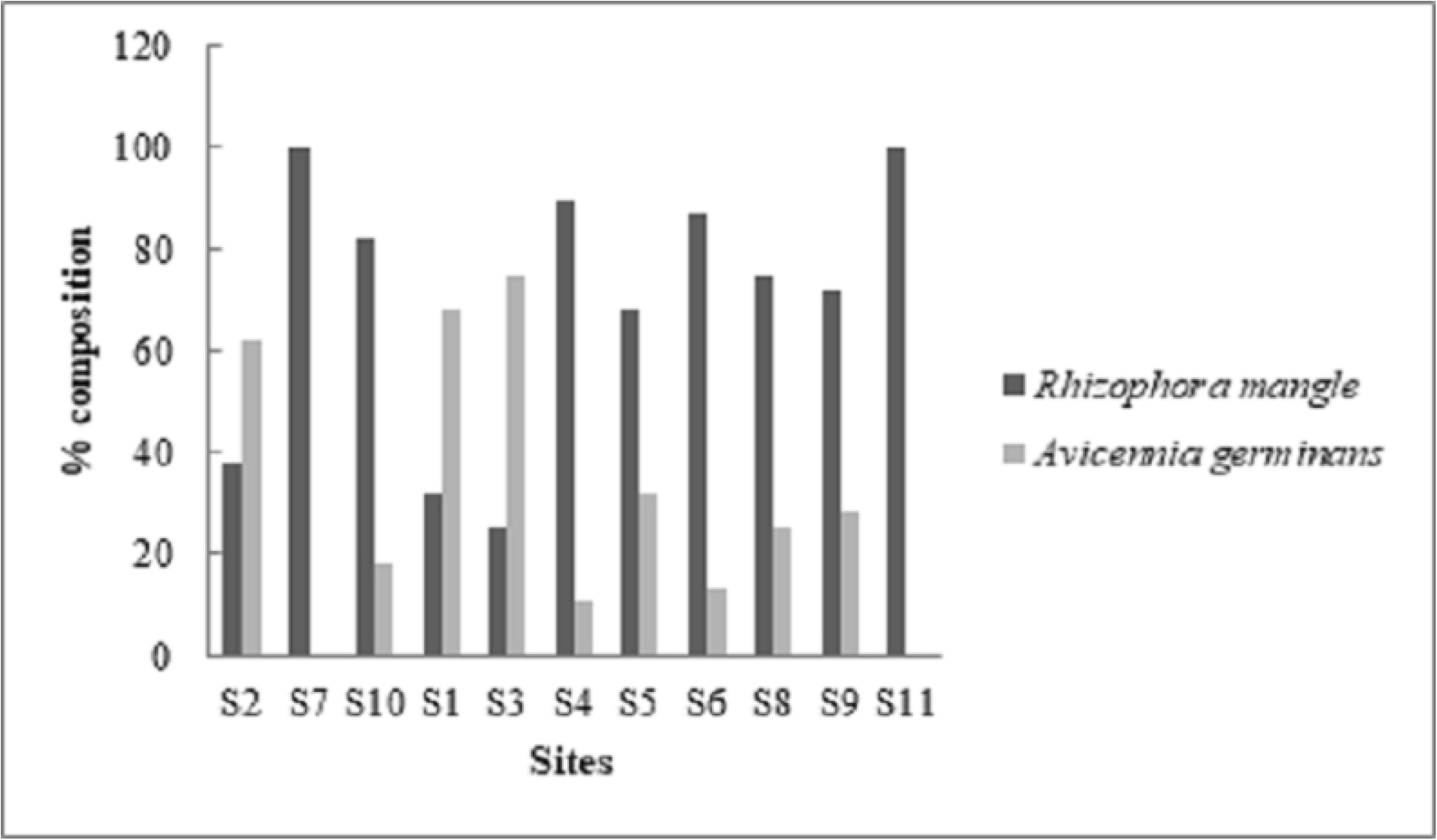
Species composition in conservation and general use zones.

**Fig 3.**
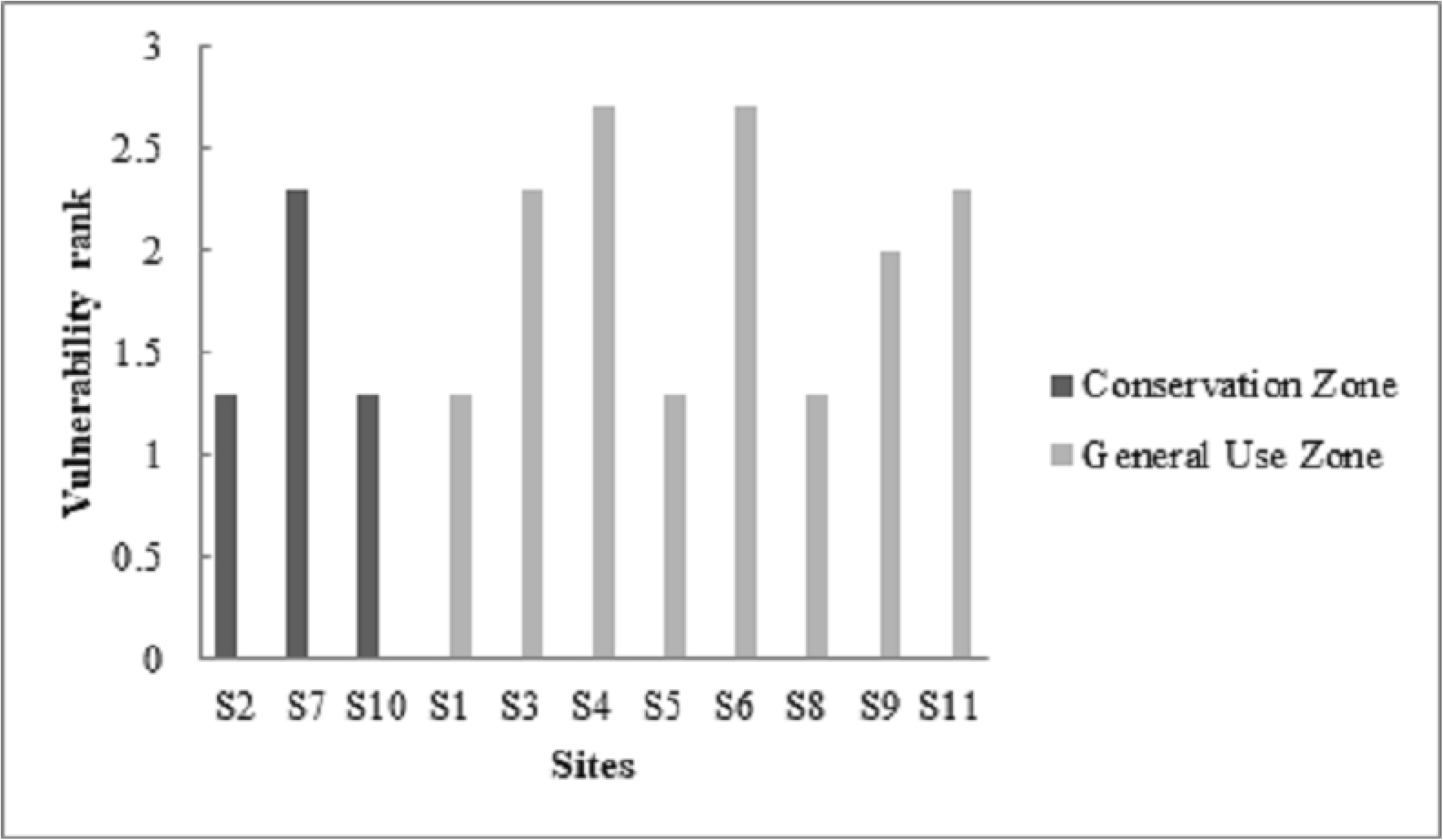
Comparison of mangrove vulnerability ranks in conservation and general use zones.

**Fig 4.**
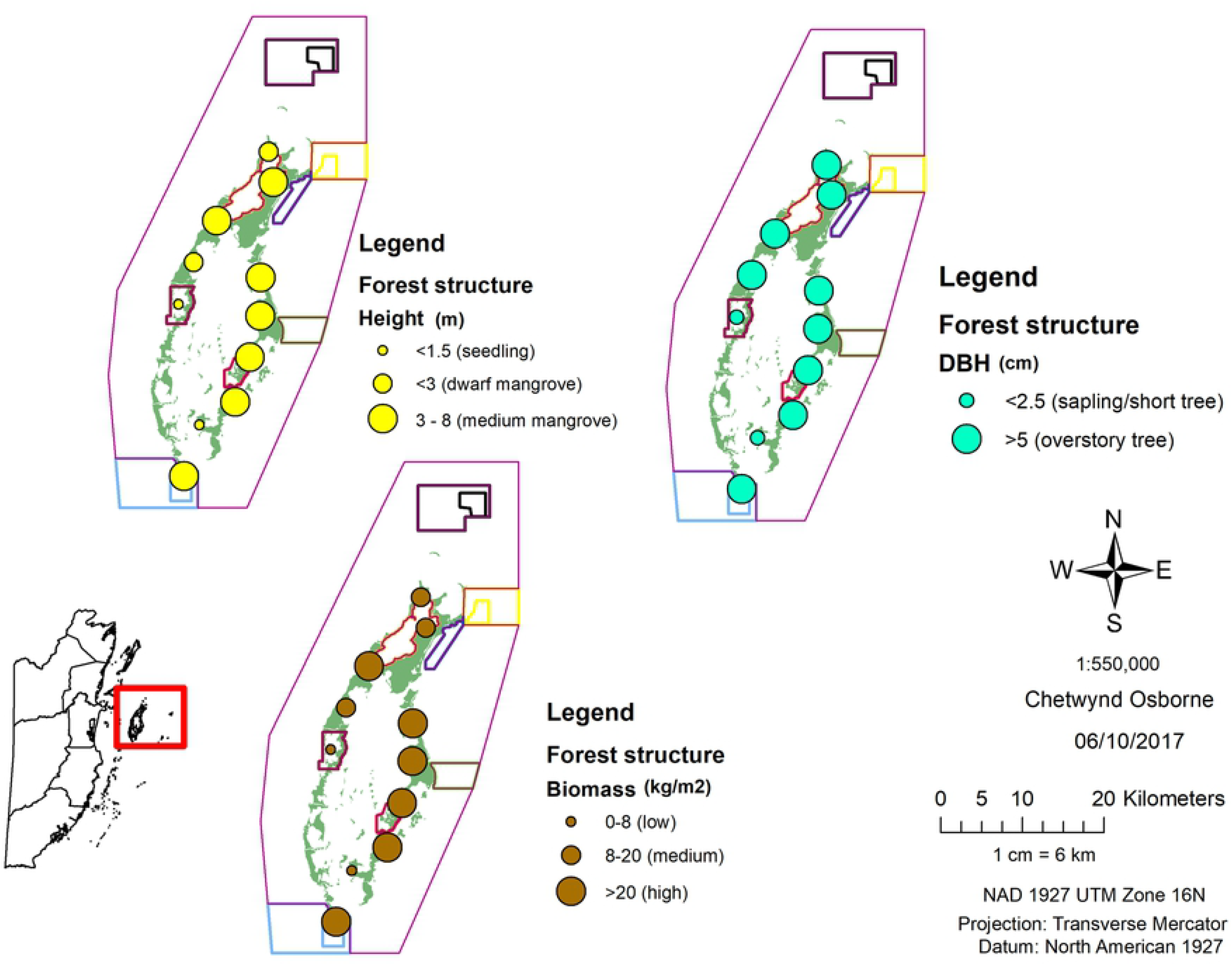
Mangrove forest structure across sites surveyed based on classification scheme of [12,16,19,20,67].

### Comparison of mangrove resilience in conservation and general use zones

Taking all trees into account (Table 3), it was evident that site two (conservation zone V) and site eight (general use zone, close to central lagoon) had higher mean height and DBH, suggesting that these were the most mature mangrove stands among surveyed sites. Site seven (preservation zone VII) and site 11 (general use zone, in central lagoon) had lower mean height and DBH (Table 3), suggesting the least mature mangrove stands among surveyed sites and these sites were densely populated with *R. mangle* seedlings. The mean height of trees for sites one, five, nine and 10 seemed to be similar (Table 3), possibly indicating similar maturity among these sites. No statistically significant difference in tree heights (Mann–Whitney test, *p* = 0.32) and DBH (Mann–Whitney test, *p* = 0.76) existed between sites in conservation and general use zones. Based on mean height and DBH, tall mangrove stands were absent among surveyed sites. Dwarf mangroves and mangrove seedlings were the least extensive, while medium mangroves were the most extensive, found at sites in conservation and general use zones (Table 3). Mangrove standing biomass ranged from 0.11 to 95.91 kg/m^2^ among sites surveyed (Fig 5) and site eight (general use zone, close to central lagoon) had the highest standing biomass while site seven (preservation zone VII) had the lowest. Other sites in the general use zone (such as sites one, five, and nine) and conservation zone (such as sites two and 10) had similar standing biomass (47.30 ± 6.13 kg/m^2^). No statistically significant relationship (*p* = 0.71, Spearman’s correlation) existed between *R. mangle* prop root surface area and *R. mangle* standing biomass, indicating that *R. mangle* prop root surface area was not strongly dependent on *R. mangle* standing biomass.

**Table 3.** Summary of sites based on mangrove type classification scheme of [12] and [16].

| Sites | Zones | Mean DBH ± SD (cm) | Mean height ± SD (m) | Mangrove type |
| --- | --- | --- | --- | --- |
| S1 | General use | 26.22 ± 15.25 | 4.48 ± 2.14 | Medium |
| S2 | Conservation | 30.30± 15.30 | 4.99 ± 1.97 | Medium |
| S3 | General use | 20.96 ± 18.81 | 3.46 ± 2.35 | Medium |
| S4 | General use | 13.61 ± 7.97 | 2.90 ± 0.80 | Dwarf |
| S5 | General use | 24.62 ± 15.63 | 4.20 ± 2.03 | Medium |
| S6 | General use | 14.17 ± 10.29 | 2.26 ± 0.94 | Dwarf |
| S7 | Conservation | 3.02 ± 4.82 | 1.23 ± 0.42 | Seedling |
| S8 | General use | 32.61 ± 13.00 | 6.44 ± 1.81 | Medium |
| S9 | General use | 26.08 ± 18.09 | 4.64 ± 1.90 | Medium |
| S10 | Conservation | 27.59 ± 15.02 | 4.15 ± 1.81 | Medium |
| S11 | General use | 1.49 ± 0.44 | 0.64 ± 0.11 | Seedling |

**Fig 5.**
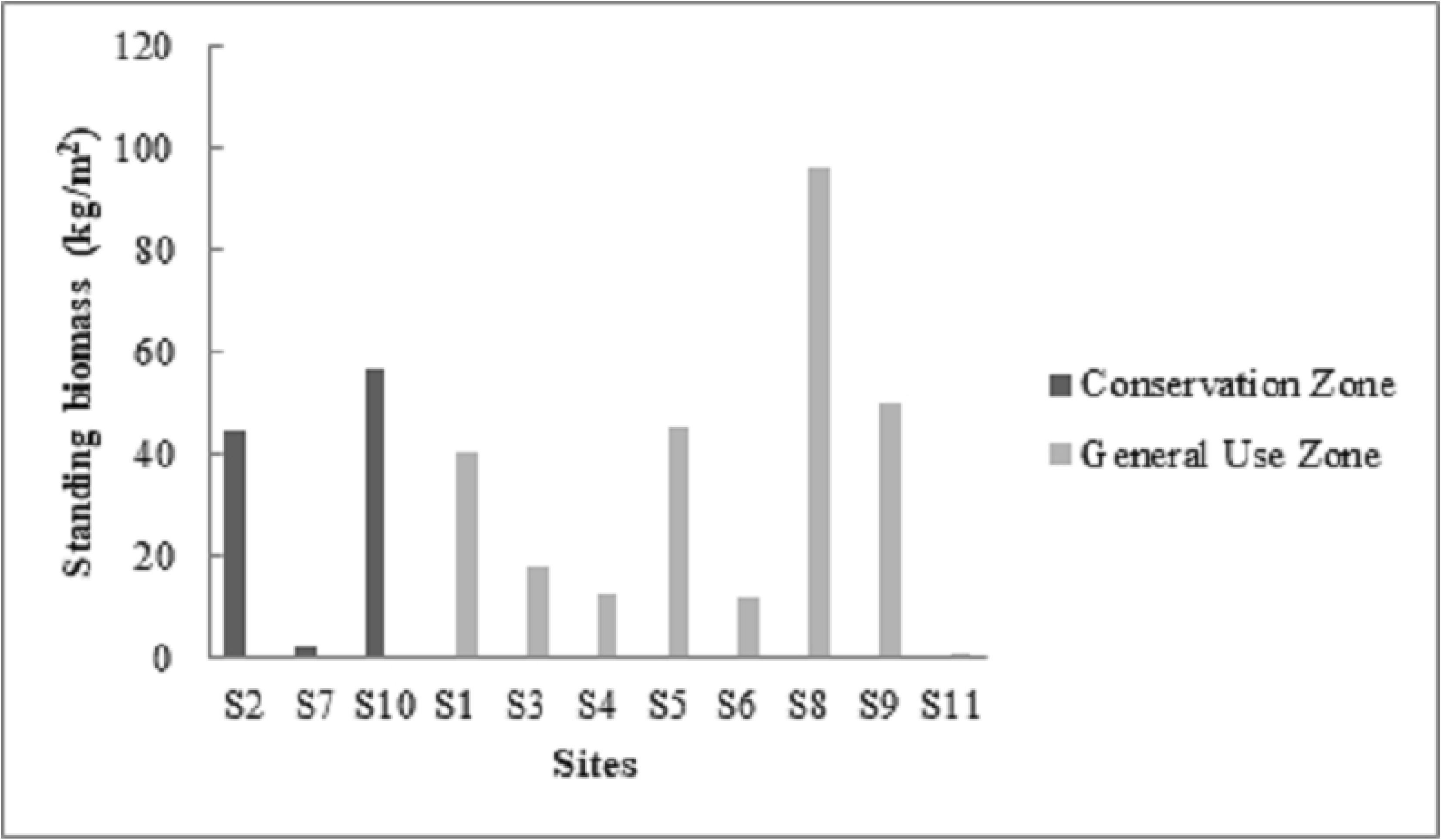
Comparison of mangrove standing biomass in conservation and general use zones.

### Effect of mangrove prop root structure on fish populations

The sites surveyed had a fairly rich diversity of fishes belonging to 17 families and 35 species. The abundance and size ranges of fish by the dominant families (Haemulidae, Lutjanidae, Pomacentridae, and Scaridae) were used to determine the nursery function of mangrove on fish populations and related to mangrove health or resilience. Site one (general use zone, in central lagoon) had the highest Shannon-Weiner diversity index value (H^’^ = 2.17) while site eight (general use zone, close to central lagoon) had the lowest Shannon-Weiner diversity index value (H^’^ = 0.18) (Table 5). Diverse communities also existed for other sites in conservation (S2, S7, and S10) and general use (S4, S9, and S11) zones based on Shannon-Weiner diversity index values (Table 5).

**Table 5.**
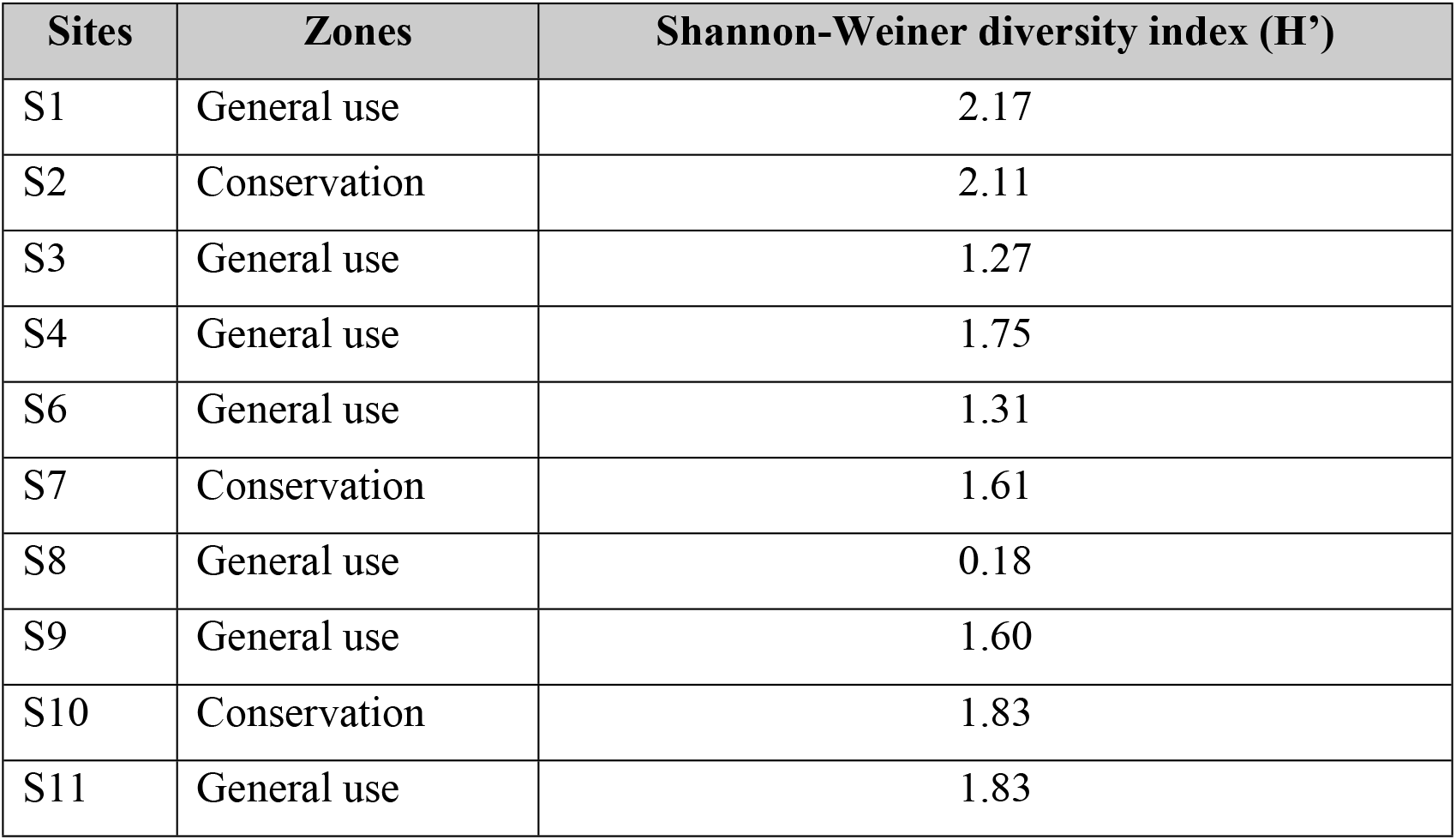
Diversity of ichthyofauna in conservation and general use zones.

| Sites | Zones | Shannon-Weiner diversity index (H') |
| --- | --- | --- |
| S1 | General use | 2.17 |
| S2 | Conservation | 2.11 |
| S3 | General use | 1.27 |
| S4 | General use | 1.75 |
| S6 | General use | 1.31 |
| S7 | Conservation | 1.61 |
| S8 | General use | 0.18 |
| S9 | General use | 1.60 |
| S10 | Conservation | 1.83 |
| S11 | General use | 1.83 |

The results of mangrove prop root measurements in sub-tidal zones are shown in Fig 6, where higher prop root measurements were recorded for the conservation zone. Statistically significant relationship (*p* = 0.01, Spearman’s correlation) existed between fish abundance and prop root surface area. Haemulidae and Lutjanidae were the most abundant families and high fish abundance was recorded for sites one (general use zone, in central lagoon), two (conservation zone V), seven (preservation zone VII) and 10 (conservation zone VI) (Fig 7). Pomacentridae and Scaridae were the least abundant families and low abundance was recorded for sites five (general use zone, close to Vincent’s lagoon special management area) and 11 (general use zone, in central lagoon) (Fig 7). Based on total fish biomass for conservation and general use zones (Fig 8), Lutjanidae and Haemulidae had high total fish biomass while Pomacentridae and Scaridae had low total fish biomass. Sites seven and 10 had the highest total fish biomass (Fig 8). Although site one had high fish abundance, the total fish biomass was low (Fig 8), suggesting the presence of very small fishes. From Fig 9 it was evident that the highest density was recorded for Haemulidae and Lutjanidae within size class 6-10 cm while the lowest density was recorded within size class 31-40 cm, and all families had fairly representative densities within size class 0-5 cm. Size classes of 0-5 cm and 6-10 cm were most frequent among certain general use zone sites (such as S1, S3, S4 and S11) and conservation zone site (such as S2). Size classes of 11-20 cm, 21-30 cm and 31-40 cm were more frequent among sites seven and 10. No statistically significant difference (*p*>0.05, *t*-test) was found in estimation of fish abundance between study sites in conservation and general use zones. Prop root surface area was isolated by multiple linear regression (*p*=0.02) as the main variable influencing fish species assemblage.

**Fig 6.**
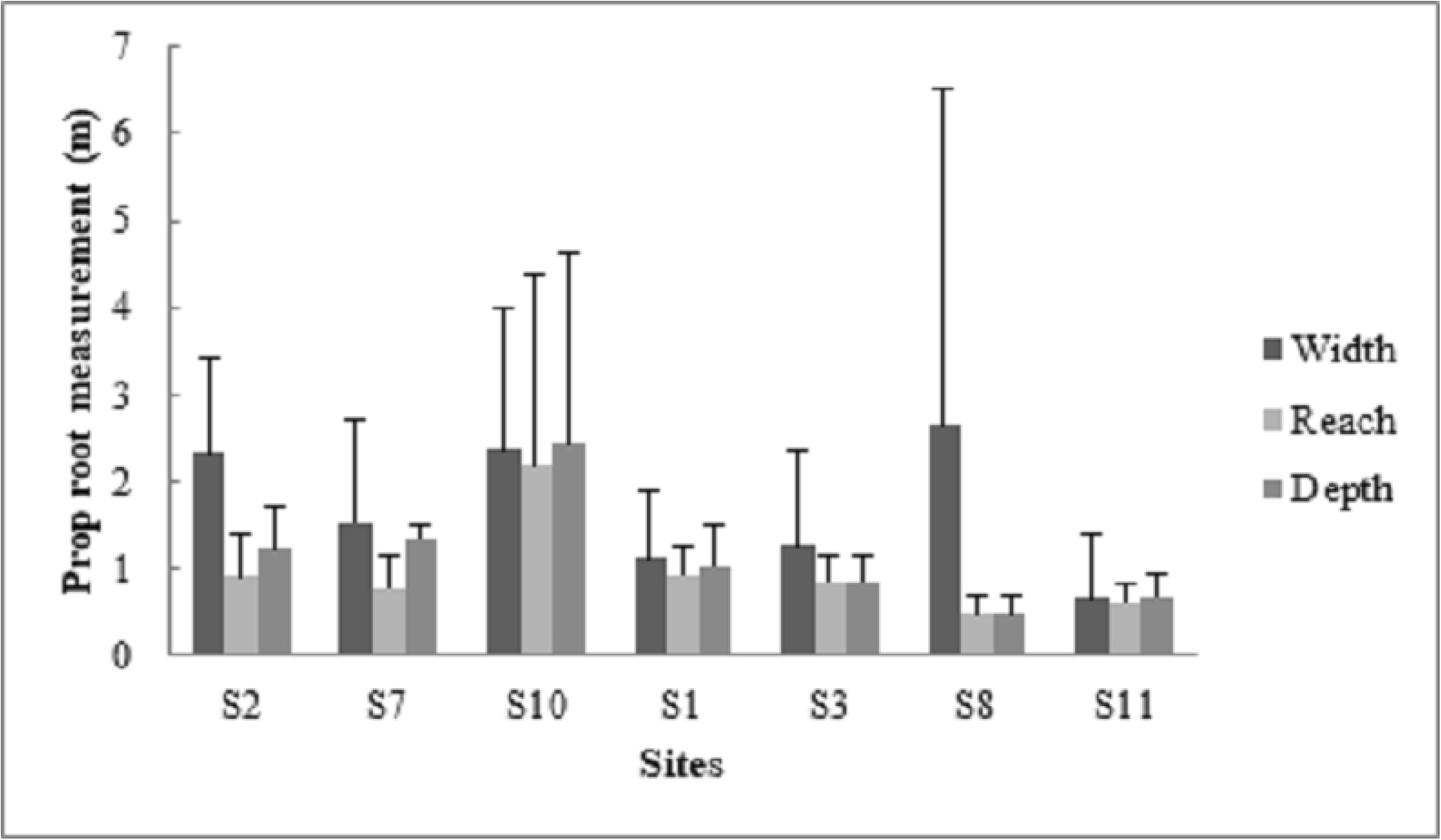
Sub-tidal measurements of prop root zones across sites.

**Fig 7.**
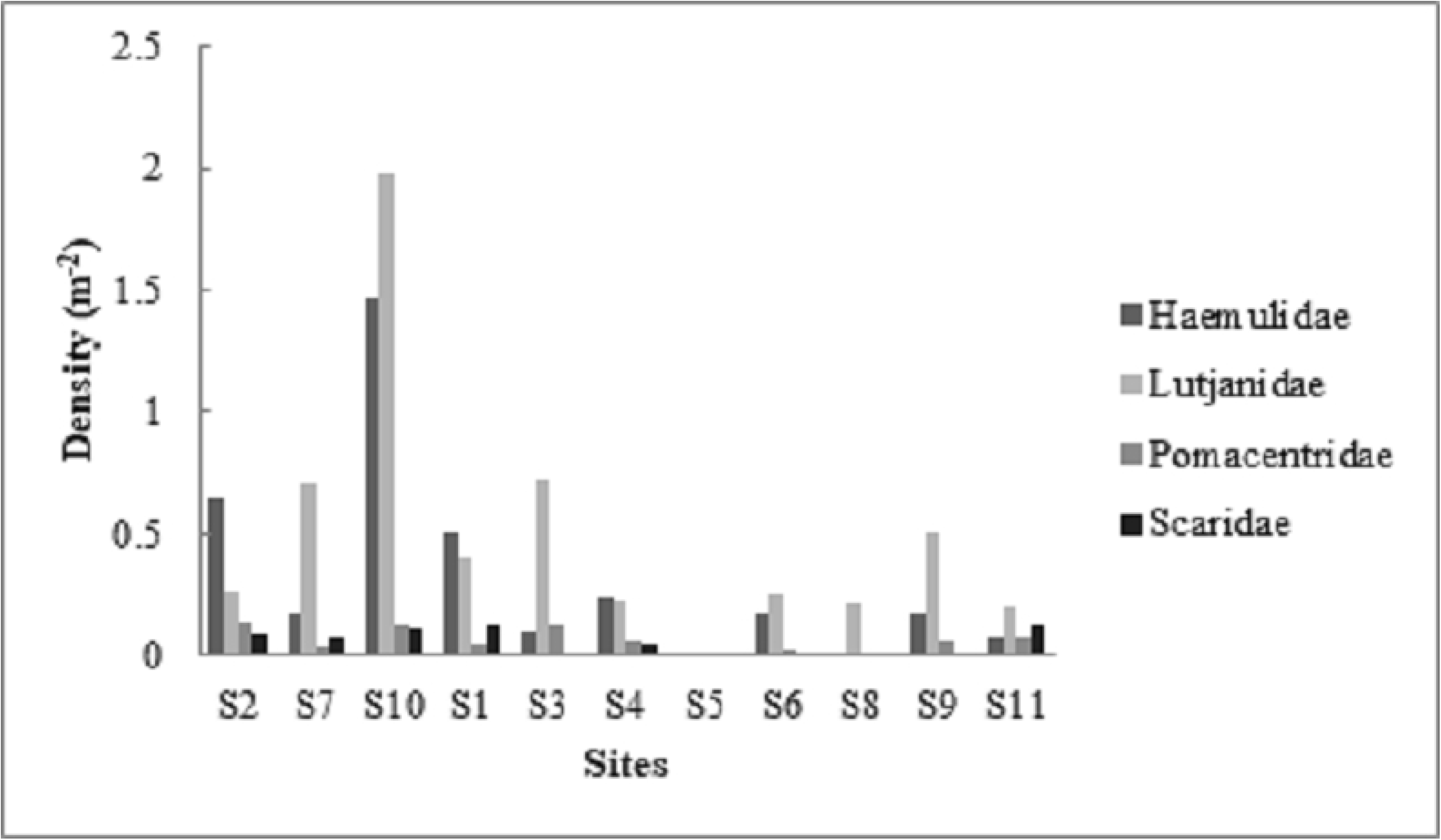
Comparison of fish abundance across surveyed sites.

**Fig 8.**
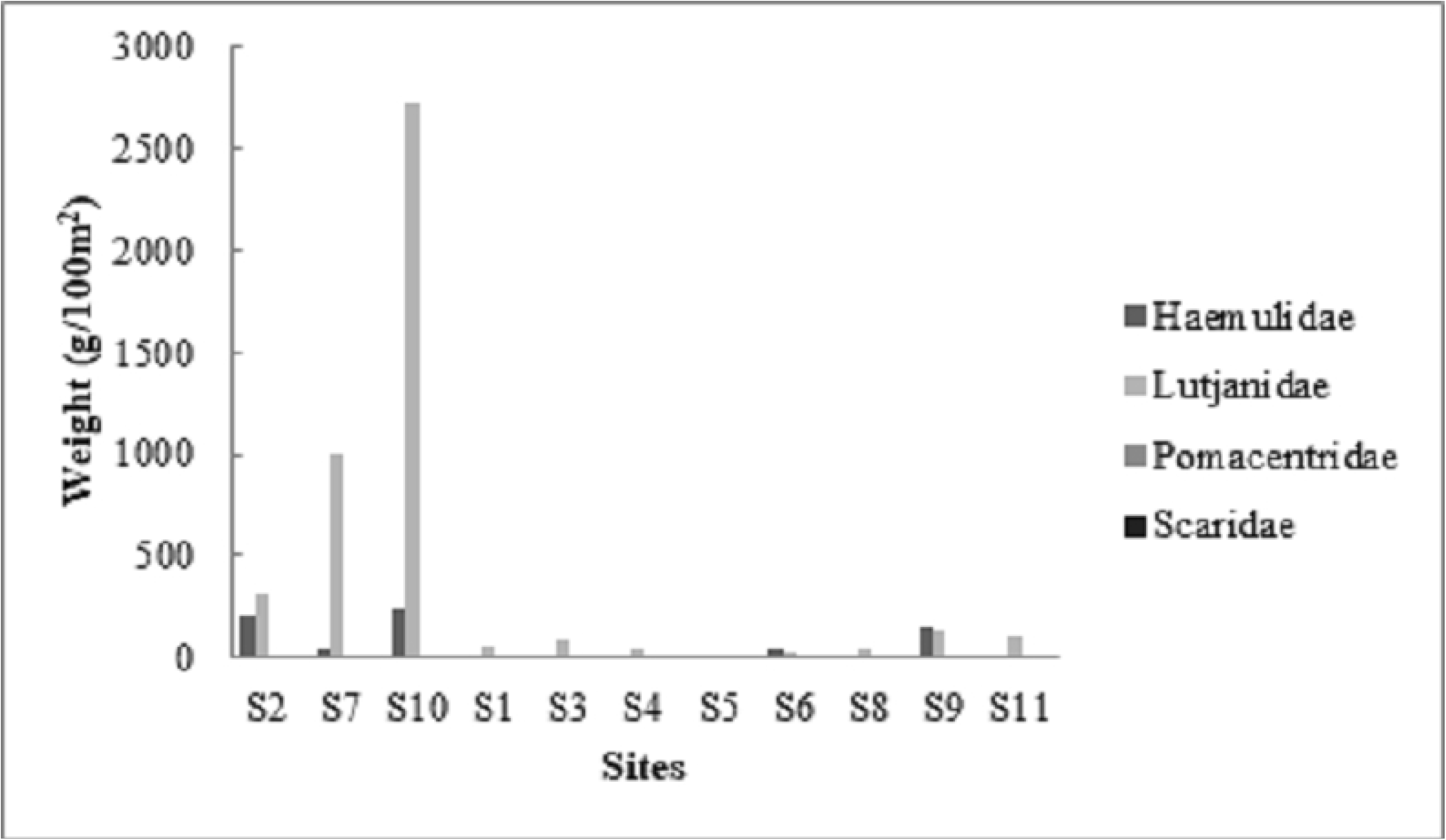
Comparison of total fish biomass across surveyed sites.

**Fig 9.**
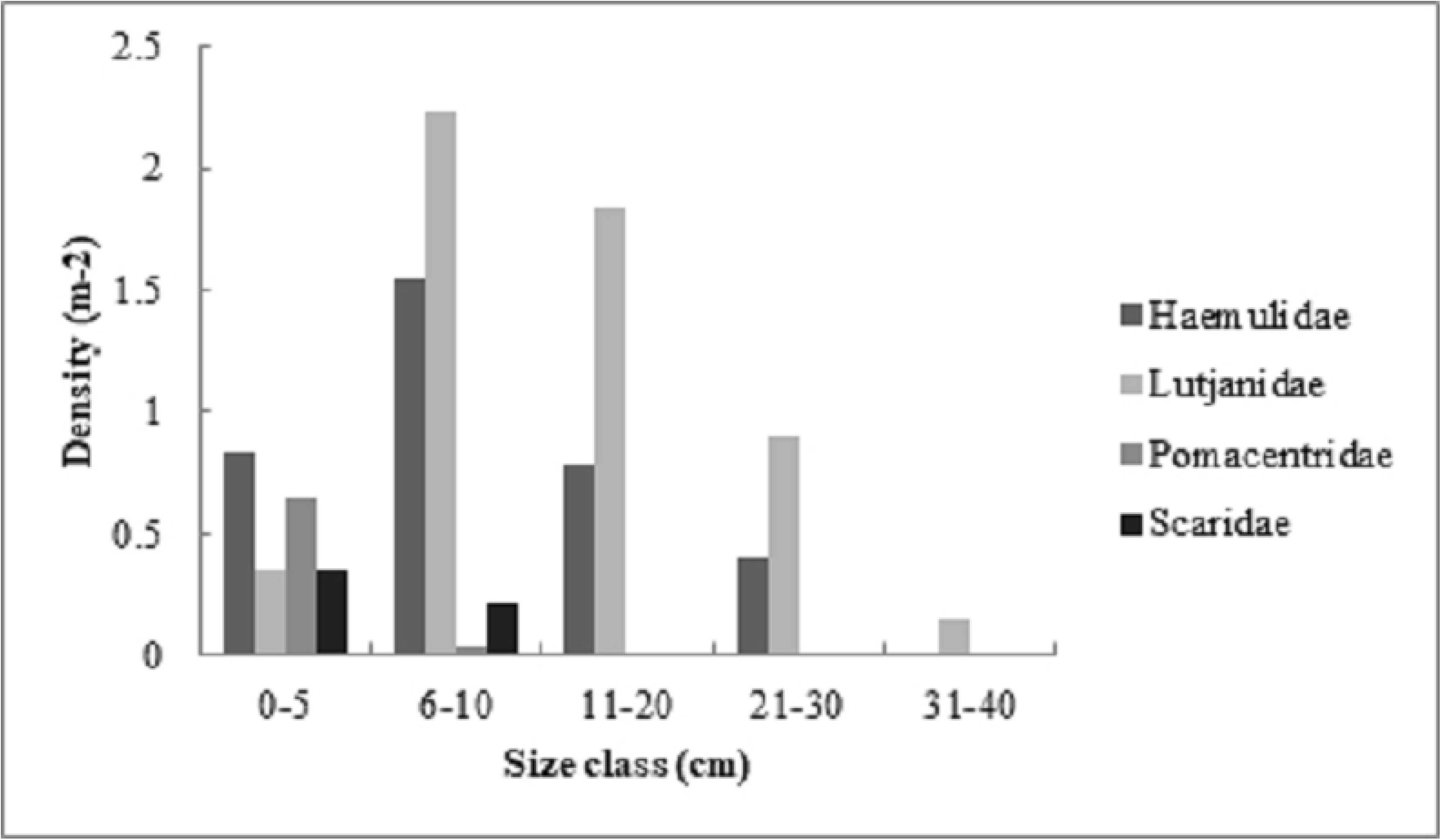
Comparison of fish density by size class across surveyed sites.

## Discussion

The dominance of *R. mangle* across surveyed sites suggested that land-building is taking place [68]. The dominance of *A. germinans* for sites one, two, and three may be attributed to the close location of these sites to sheltered areas such as lagoons which provides protection from direct exposure to the striking of large waves found on the outer atoll since *A. germinans* lack the network of prop roots present in *R. mangle* to disintegrate wave action [69,70]. Results of the vulnerability assessment ranking found that all sites had some components of vulnerability. Sites S1, S2, S5, S8, S9, and S10, with ranking of 1 – 2, indicated mangrove areas with current resilience and mangrove resilience could enhance with rank reduction of any vulnerability components higher than 4 [17]. Ranks of 2 – 4 (such as S3, S4, S6, S7, and S11) indicated mangrove areas that were prone to some core vulnerability which could be improved by targeted management [17]. These results further emphasize the need for the establishment of more mangrove conservation areas that will facilitate sustainable use of resources to better control non-climate stressors and protect mangrove areas in the general use zone. This sort of control and protection can be achieved through improved local management and reduction of human impacts [17,50], which will increase resilience of species and mangrove habitats to the effects of climate change [17,71]. Some local management strategies that can be employed include sustainable use of available resources to maintain and enhance ecosystem functions [72], the utilization of inclusive governance structures at the community level to balance the exploitation and conservation of mangrove ecosystems [72,73] and measures to control land reclamation, boat anchoring, sediment dredging, and coastal and aquaculture development [74]. Sites in conservation zones had a small amount of rubbish as compared to some sites in the general use zone which had large amounts of rubbish. This sort of human impact may have contributed to the inherent vulnerability in the general use zone. Additionally, rubbish accumulation may contribute to the establishment of dry land [68,75] which was evident in some transect areas and this sort of dry and bare composition of soil may account for the lack of mangroves in some transect areas.

Sites in conservation and general use zones were all a part of Turneffe Atoll; hence, similarities among the structure of mangrove communities present at these sites. A phenomenon of this nature was evident since no significant difference existed for tree heights and DBH between sites in conservation and general use zones. The fairly wide distribution of DBH and high DBH measurements among sites in conservation (S2 and S10) and general use (S1, S5, S8, and S9) zones, indicated uneven-aged and more mature and healthier mangrove stands [20]. Additionally, the extensive distribution of overstory trees (DBH > 5 cm) across surveyed sites provided more corroborating evidence about the extensive cover of old mangrove stands [21] in Turneffe Atoll. The nearly continuous barrier reef that runs along the coastline of Belize provides shelter for the shore, absorb majority of the inward wave energy, and provides suitable conditions for the establishment of mangrove seedlings [12]. A situation of this nature may in part account for the dense population of *R. mangle* seedlings present in preservation zone (site seven) and general use zone (site 11, in central lagoon). This large proportion of seedlings, indicates colonization of forest gaps [21] among these sites.

The results of mean DBH and height across surveyed sites showed that dwarf mangroves (typically < 3 m tall and DBH > 5cm) were among the least extensive mangrove type represented across surveyed sites. Therefore, majority of the sites surveyed in Turneffe Atoll lacked ideal environmental conditions which restrict mangrove vegetation to a scattered cover of short mangroves. The presence of dwarf mangroves in certain sites indicated mangrove resilience since dwarf mangroves can survive natural disturbances such as hurricanes which are accompanied by high water levels that cover these small individuals, preventing strong winds from blowing them down [4]. This sort of survivability of dwarf mangroves renders them as possible sources of propagules for recolonization processes subsequent to major disturbances [4]. Medium mangroves (approximately 3 – 8 m tall) were the most extensive type represented across surveyed sites, indicating that majority of the sites surveyed were densely populated by fringing *R. mangle* growing at exposed sites along the coast and on cays [12]. The site with the highest standing biomass (site eight, general use zone) can possibly be attributed to being a more productive and older mangrove ecosystem with higher total basal area [76] and more resilience as compared to the site with the lowest standing biomass (site seven, preservation zone VII). It is expected that the preservation zone should have good mangroves since the primary objective of this zone is to preserve an entirely natural state [49]. However, this preservation zone site had the lowest standing biomass and was constituted mostly by young individuals with low DBH, which is a predictive variable that is frequently used for mangrove forest standing biomass [76]. Site seven had good recruitment levels since all species were producing seedlings, indicating some amount of resilience and regeneration capabilities [17,77]. This sort of regeneration aids in the establishment and maintenance of mangrove species zonation patterns [78].

Fish measurements showed difference in species diversity among sites based on Shannon-Weiner indices (H’), where high H’ value was an indication of a more diverse community and even distribution of species abundance among all species recorded for that community [79]. Commercial fishing is allowed in the general use zone, but not in conservation zones since a no-take regime is maintained here [49]. Therefore, it is expected that fish composition in the general use zone differs from that of conservation zones; however, this was not the case since no statistically significant difference existed for fish families composition among conservation and general use zones based on t-test (*p* = 0.28). Water clarity for visual census was poor for site five and this limitation may have contributed to the low diversity and abundance recorded for this site, since visibility factors may influence the accuracy of visual censuses [7]. Based on multiple linear regression analysis, prop root surface area was the main variable influencing fish species assemblage and a strong positive correlation (r = 0.93) existed between fish abundance and prop root surface area. This strong correlation may in part account for the high fish abundance recorded among certain surveyed sites due to the fairly representative composition of *R. mangle* with large prop root structures to support fish populations. However, low fish abundance can possibly be attributed to the least mature mangrove forests dominated by *R. mangle* seedlings. Higher fish abundances were recorded for sites with medium mangrove stands (S2 and S10) as compared to other sites with seedling and dwarf mangrove stands (S11, and S4). This was possibly due to the greater shade created by medium mangrove stands [55]. These results are consistent with studies conducted by [80] and [7], which provided more corroborating evidence of the probable positive correlation of fish abundance to shade (height of trees) and increased prop roots complexity. Additionally, medium mangrove stands may have larger fringing *R. mangle* prop roots to create extensive borders around submerged habitats [81] that serve as nursery areas to support large fish populations; hence greater productivity. The high abundance of commercially important fish families such as Haemulidae and Lutjanidae for site 10 may be attributed to greater protection and reduced impact since a no-take regime is maintained for this conservation zone site as compared to sites in the general use zone where commercial fishing is allowed [49]. Pomacentridae and Scaridae are predominantly coral reef fishes that undertake ontogenetic migrations between coral reefs, seagrass beds, and mangroves [81] and the unique nature of Turneffe Atoll in terms of high connectivity among mangrove stands, seagrass, and coral reefs [49] creates an ideal opportunity for migration. Therefore, the low fish abundance recorded across study sites for Pomacentridae and Scaridae may be attributed to the seldom utilization of mangrove habitats for food and shelter. Fish species belonging to the family Scaridae are protected by legislated regulations under the Fisheries Department in Belize [49], but these species are illegally fished to support fishermen livelihoods [82]. This sort of illegal fishing may in part account for the low abundance of Scaridae across surveyed sites. The smaller mode size range frequency (0-5 cm and 6-10 cm) for sites such as one, two, and three may account for the low total fish biomass recorded for these sites, even though high fish abundance was recorded here. Additionally, the high total fish biomass for sites seven and 10 may be attributed to the larger mode size range frequency (11-20 cm and 21-30 cm). The representative fish abundance and biomass recorded across surveyed sites shared some similarities with small scale experimental studies conducted by [83] on recruitment and juvenile reef fish abundance in the Caribbean, which found higher abundance among mangrove stands since these habitats are further away from the main reef, hence reduced predation. The high fish biomass recorded for commercially important families such as Haemulidae and Lutjanidae [84] gives an indication of mangrove resilience to support these species. Moreover, the high connectivity of mangrove stands to reef and seagrass at Turneffe Atoll, provides ideal nursery habitats for commercially important juvenile fishes [49]. Therefore, the need exists for the establishment of more mangrove conservation areas in Turneffe Atoll to prolong these vital ecological goods and services that the Belizean population depends upon for their livelihoods.

Based on fish size class measurements, more nursery habitats were provided by *R. mangle* prop root structures across sites in conservation and general use zones for fish populations. The mixture of fish densities recorded for all size classes were consistent with evidence that some juvenile fishes primarily utilize mangroves for shelter while feeding opportunistically [55]. Additionally, this research was conducted during the wet season in Belize and this season tends to correlate with high recruitment of juvenile fishes to mangrove communities since a high abundance of zooplankton exists to provide greater food abundance [33,85,86]. A phenomenon of this nature provides further evidence for the high abundance of juvenile fishes within mangrove stands.

Based on the results of this study more resilient and healthier mangrove stands constituted sites (especially sites one, two, eight and 10) in the conservation and general use zones, of which the general use zone sites are not well protected in Turneffe Atoll Marine Reserve (TAMR). Based on the sample design for this study, surveyed sites were selected based of mangrove distribution and only three conservation zone sites had a representative distribution of mangroves as compared to eight sites in the general use zone. Therefore, mangroves in conservation zones may be underrepresented and the establishment of more conservation areas in the general use zone will aid in the protection of more resilient mangroves given the importance of setting conservation priorities based on how represented mangroves are in current conservation zones [87]. This sort of protection is critical given the connectivity that mangroves share with other marine ecosystems to foster higher immigration rates from nursery habitats and possibly more production to support livelihoods [81]. Site eight is close to Blackbird Caye conservation zone which is privately owned [88] and slated for further development given the major role that tourism plays in the economic development of Belize [84]. This sort of development will impact mangrove ecosystems and lead to irreversible losses of mangroves, thereby threatening mangrove resiliency [44]. Mangrove losses can also affect water quality since pollutants from waste water would not be readily removed and transformed [89,90] and fish abundance may decline since nursery habitats which provide shelter for juvenile fishes are removed [91]. Therefore, more careful evaluation of conservation measures is needed for mangroves specifically. Some of these conservation measures include: (a) greater elimination of non-climate stressors on mangroves such as pollution, filling, dredging, land conversion for aquaculture in order to improve ecosystem health to some extent and minimize mangroves vulnerability to and increase their resilience to climate change stressors [92,93]; (b) establishment of more mangrove conservation areas where no mangrove clearing is allowed will prevent destruction to mangrove nursery areas, thereby reducing the impacts on other nearshore marine ecosystems (*e.g*., coral reefs) [94]. These conservation areas can predominantly target the protection of mangroves in the general use zone and allow commercial fishing in a sustainable manner since the general use zone provides fertile and valuable fishing ground; and (c) implementation of more outreach and educational programmes in order to garner greater community support and solidify actions that will aid in enhancing mangrove ecosystems resilience and resistance to climate change [93]. Additionally, these outreach and educational programmes will improve our understanding of the drivers and value of mangrove ecosystem services, which may further boost efforts to better manage mangroves [67]. Combining visual censuses with unattended, continuous-recording method such as underwater video system would answer additional questions [94].

## Conclusion

The physical and ecological characteristics of Belize are similar to other parts of the Caribbean and Central America. However, in the region Belize is unusual since a large portion of the coastline is covered by mangroves and an extensive composition of mangroves is found at Turneffe Atoll. Fringing *R. mangle* is vital for Belize due to the region’s low tidal range which allows mangrove prop root to remain permanently inundated, providing a predictable nursery habitat for juvenile fishes. This sort of mangrove characteristics is proof of mangrove resilience within Turneffe Atoll. Mangroves in conservation zones at Turneffe Atoll are possibly underrepresented since a larger composition of resilient mangroves constitutes the general use zone. Major gaps exist in the protection of coastal and marine mangroves within Turneffe Atoll and although the findings of this research only form a baseline, the need exists for the establishment of more mangrove conservation areas in Turneffe Atoll due to the presence of resilient and healthy mangroves in the general use zone. The establishment of more mangrove conservation areas would better facilitate more effective management of fisheries, shoreline protection and increase resilience to climate change impacts.

## Acknowledgements

Gratitude is extended to the following organizations that provided financial support for this research: CARPIMS and The University of Belize Environmental Research Institute. Thanks to the staff and personnel of University of Belize Calabash Caye Field Station, for their hospitality, provision of research materials, and logistical support. Thanks to Tracy Gazobatu, Elisabeta Maiwiriwiri, and Roberto Gongora for assistance with the field work. The comments of two anonymous reviewers significantly benefited this paper.

